# Requirement for Anti-Apoptotic MCL-1 during Early Erythropoiesis

**DOI:** 10.1101/2020.05.06.081422

**Authors:** Meghan E. Turnis, Ewa Kaminska, Kaitlyn H. Smith, Brittany J. Kartchner, Peter Vogel, Jonathan D. Laxton, Richard A. Ashmun, Paul A. Ney, Joseph T. Opferman

## Abstract

Mature erythrocytes are under tight homeostatic control with the need for constant replacement from progenitors to replace damaged or obsolete red blood cells (RBCs). This process is regulated largely by erythropoietin (Epo) which promotes the survival of erythroid progenitors and facilitates their differentiation and proliferation. Ablation of *Bcl2l1* (which encodes BCL-xL) results in embryonic lethality with a lack of mature erythrocytes but does not perturb erythroid progenitors. Similarly, conditional *Bcl2l1*-deletion results in severe anemia with the death of late erythroid progenitors and induction of extramedullary erythropoiesis. While BCL-xL is critical to the survival of mature erythrocytes, it is still unclear whether other anti-apoptotic molecules mediate survival during earlier stages of erythropoiesis. Here, we demonstrate that erythroid-specific *Mcl1*-deletion results in embryonic lethality due to severe anemia caused by a lack of mature RBCs. *Mcl1*-deleted embryos exhibit stunted growth, ischemic necrosis, and decreased RBCs in the blood. Furthermore, we demonstrate that the dependence on MCL-1 is only during early erythropoiesis, whereas during later stages the cells become MCL-1-independent and upregulate the expression of BCL-xL. Functionally, MCL-1 relies upon its ability to prevent apoptosis to promote erythroid development since co-deletion of the pro-apoptotic effectors *Bax* and *Bak* can overcome the requirement for MCL-1 expression. Furthermore, ectopic expression of human BCL2 in erythroid progenitors can compensate for *Mcl1* deletion, indicating redundancy between these two anti-apoptotic family members. These data clearly demonstrate a requirement for MCL-1 in promoting survival of early erythroid progenitors.

## Introduction

Erythropoiesis requires the cytokine erythropoietin (Epo), which promotes survival during red blood cell (RBC) development and facilitates the differentiation of erythroid progenitors^1,2^. Binding of Epo to its receptor (EpoR) induces activation of JAK2, and recruitment of STAT5a/b^3,4^. BCL-xL expression can be induced during terminal erythroid differentiation^5,6^ and STAT5 binds to the *Bcl2l1* (encodes BCL-xL) promoter^7^. However, there is controversy about the requirement for STAT5a/b during erythropoiesis and whether it is needed to induce BCL-xL in response to Epo^8^. Furthermore, it is still unclear whether BCL-xL is the essential survival factor downstream of Epo signaling during early erythropoiesis.

A critical feature of Epo-mediated signal transduction is promotion of erythroid progenitor survival^9^. *Epo*- and *EpoR*-deficient animals die around embryonic day 12.5 (E12.5) due to a deficiency in definitive erythropoiesis^2^. *Bcl2l1*-deficient animals (lacking BCL-xL) form blood progenitors but lack mature RBCs and die around E12.5^10^ supporting the notion that BCL-xL is a survival factor downstream of Epo signaling. However, *Bcl2l1*-conditional knockout animals survive, but develop hemolytic anemia, indicating that BCL-xL is required for promoting the survival of mature erythrocytes and not progenitors^11^. Furthermore, culture of erythroid progenitors with Epo induces BCL-xL expression 36 hours after the start of differentiation^5^. These data imply that while BCL-xL may play a critical role in mature erythrocytes, other pro-survival factors must promote the survival of erythroid progenitors.

Anti-apoptotic members of the BCL-2 family prevent the induction of apoptosis by inhibiting BAX and BAK activation^12^. Genetic models have provided insight into the roles of individual anti-apoptotic molecules during hematopoiesis. *Bcl2*-deficient mice are viable, but exhibit a spectrum of abnormalities including lymphocyte apoptosis, but otherwise hematopoiesis proceeds unperturbed^13-15^. Loss of all three isoforms of *Bcl2a1* causes only minor defects in hematopoiesis^16^. BCL-w is not required for hematopoiesis^17-19^. BCL-xL is required for late erythropoiesis and protection of mature platelets^10-20,21^. In contrast, *Mcl1* is essential for survival of multiple hematopoietic lineages including stem cells^22^, B cells^23-25^, T cells^23-26,27^ and neutrophils^28,29^. However, whether *Mcl1* plays any role in erythropoiesis is unknown.

Here, we report for the first time that conditional *Mcl1*-deletion, using an erythroid-specific mouse which contains the *GFPcre* gene knocked-in to the Epo receptor (*EpoR*) locus30, leads to failure of RBC maturation, anemia, and embryonic lethality. Using an *ex vivo* culture system, we show that expression of MCL-1 is only required during early erythropoiesis, but is dispensable later. Finally, *Bax*- and *Bα*/-deletion, or lentiviral-mediated overexpression of the pro-survival factor *BCL2* can rescue *Mcl1* loss in murine erythropoiesis. These data reveal a requirement for MCL-1 during early erythropoiesis, mediated by its anti-apoptotic function.

## Materials and Methods

### Mice

*Mcl1^F/F^* and *Mcl1^F/F^* Rosa26-ERCreT2 mice are previously described^31^. EpoR-Cre^30^ mice were crossed to *Mcl1^F/F^* animals. *Bax*^F/F^*Bak*^−/−^ mice^32^ were crossed to Rosa26-ERCreT2 animals. All procedures were approved by the St. Jude Institutional Animal Care and Use Committee.

### Fetal Liver Studies

E11.5-E15.5 *Mcl1^F/F^* Rosa-ERCreT2 and C57BL/6 fetal liver cells were isolated and/or cultured^33,34^. TER119-negative cells isolated using anti-TER119-biotin microbeads and magnetic columns (Miltenyi) were seeded on fibronectin-coated plates (Corning). For inducible deletion, 4-OH-tamoxifen (4-OHT, 0.5μM, Sigma) was added during the first 18 hours of culture or overnight the second day of culture. Media conditions were: Day 1, IMDM containing 15% FBS, 1% penicillin-streptomycin, 2mM L-glutamine, 10^−4^M β-mercaptoethanol, Stem Cell Factor (50ng/mL, Rockland), FLT3 Ligand (30ng/mL, Rockland), and IL-6 (20ng/mL, Rockland). Day 2, IMDM containing 15% FBS, 1% penicillin-streptomycin, 1% BSA (Stemcell), 200μg/mL holo-transferrin (Sigma), 10μg/mL human insulin (Sigma), L-glutamine, β-mercaptoethanol, and 2U/mL Erythropoietin (R&D Systems). Day 3, IMDM containing 20% FBS, 1% penicillin-streptomycin, L-glutamine, and β-mercaptoethanol.

### Lentivirus studies

Human cDNAs cloned into pHAGE-UBC-GFP-zsGreen vector were packaged in 293T cells^35^. TER119-negative fetal liver cells were infected (MOI 5) in LentiBOOST^™^ (Sirion Biotech) for 16 hours followed by 4-OHT treatment and media changes as above.

### Transplant studies

Bone marrow (BM) from donors was transplanted (i.v.) into lethally irradiated (1100 Rad) recipients (B6.SJL-Ptprca Pepcb/BoyJ, Jackson). After engraftment,recipients received tamoxifen (Sigma) to activate Cre (1mg/day for 5 consecutive days by gavage) and were monitored for 6 weeks, then harvested.

### Pathology

Embryos fixed in 10% neutral-buffered formalin (ThermoFisher Scientific) were embedded, sectioned, mounted (Superfrost Plus; Thermo Fisher Scientific), and stained with hematoxylin and eosin (H&E). Immunohistochemical (IHC) staining with anti-AHSP (Rockland) and anti-TER119 (Biosciences) were detected using DISCOVERY ChromoMap DAB (Ventana Medical Systems).

### Cytospins and Colony-forming assays

MethoCult M3434 for BFU-E or MethoCult M3534 for myeloid (StemCell Technologies) were used for colony-forming assays. Cytospins were performed using SuperfrostPlus slides (Erie Scientific) with a Cyto-Tek centrifuge (Miles Scientific) and stained with DipQuick (Jorgensen Laboratories).

### Real-time PCR, immunoblot, and flow cytometry

RNA was extracted using Trizol (Invitrogen) or RNeasy Kit (Qiagen) and reverse transcribed using SuperScript III (Life Technologies). Real-time PCR was performed using SYBR-Green (Thermo Fisher Scientific). Data acquired using a Quantstudio7 Flex real-time PCR machine (Thermo Fisher Scientific) were analyzed by ΔΔCt method with housekeeping gene (Ubiquitin) compared to unstimulated cells. Primer sequences are available by request.

Immunoblotting was performed as previously described^36^. Antibodies against: MCL-1 (Rockland), BAK, Actin (Millipore), BAX, BCL-xL, MCL-1 (Cell Signaling), BCL-2 (BD Biosciences), and anti-rabbit, -hamster, or -mouse horseradish peroxidase–conjugated secondary antibodies (Jackson Immunochemical).

Flow cytometry was acquired on LSRII, LSRFortessa, or FACSCanto II (BD Biosciences) cytometers with the Flow Cytometry and Cell Sorting Shared Resource.

Sorting was performed using Aria sorters (BD Biosciences). Data were analyzed in FlowJo V10 (FlowJo). Analyses consisted of standard two-dimensional gating or t-distributed stochastic neighbor embedding (tSNE) analysis, for 2D visualization of high-dimensional data while preserving the data structure^37^.

Statistical analyses were performed using GraphPad Prism (GraphPad Software).

## Results

### MCL-1 is expressed early in erythropoiesis

To explore the expression of anti-apoptotic family members during erythropoiesis, wild-type E12.5 fetal livers were sorted into populations correlating with their differentiation stages33. Analyses revealed a stepwise progression from S0 (primitive progenitors; TER119^−^, CD71^−^), S1 (proerythroblasts; TER119^−^, CD71^+^), S2 (early basophilic erythroblasts; TER119^low^, CD71^+^), S3 (late basophilic erythroblasts; TER119^high^, CD71^+^), S4 (chromatophilic and early orthochromatophilic erythroblasts; TER119^high^, CD71^low^), and S5 (late orthochromatophilic erythroblasts and reticulocytes; TER119^high^, CD71^−^) (Figure 1A). Quantitative PCR of mRNA from sorted populations revealed that S0 progenitors express multiple anti-apoptotic family members (MCL-1, BCL-xL, and BCL-w), but as the cells differentiate into S1/2 progenitors the predominantly expressed anti-apoptotic molecule is MCL-1 (Figure 1B). As the erythroid cells continued to differentiate into S3 and S4 stages, MCL-1 levels remained stable while BCL-xL expression increased.

**Figure 1.**
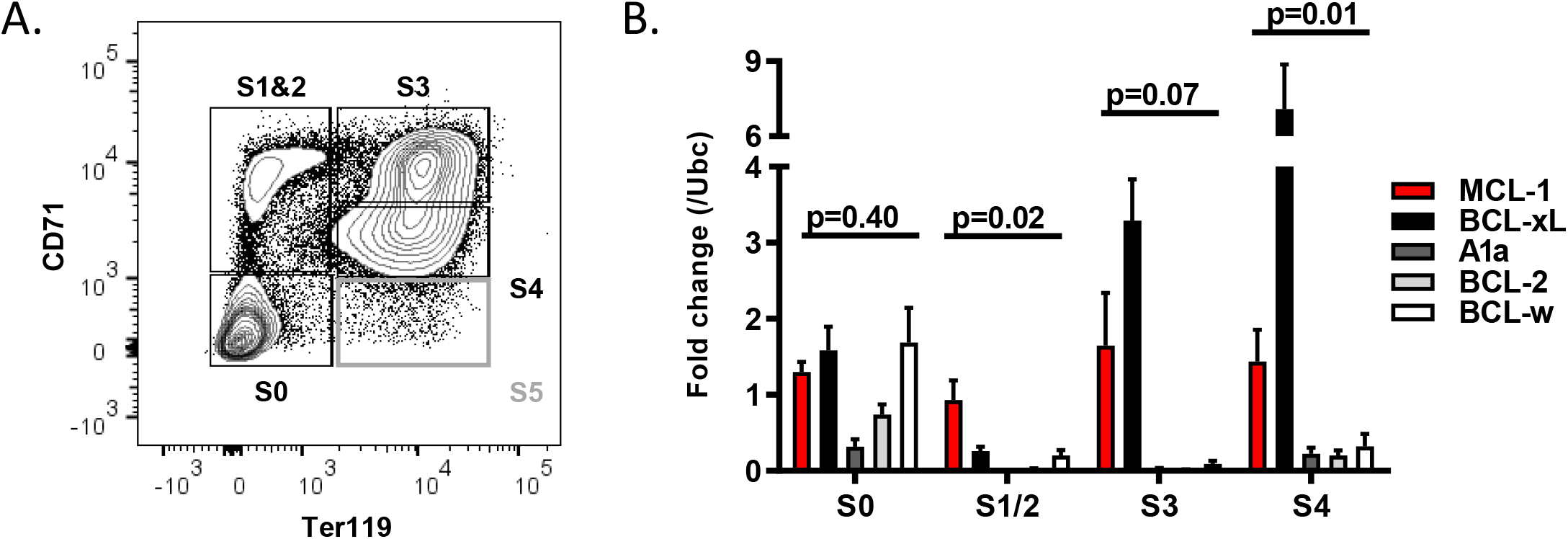
MCL-1 is expressed in early erythropoiesis. (A, B) Mouse fetal livers were isolated from E12.5 embryos and mechanically dissociated. Fetal liver cells (FLC) were labeled with CD45, CD71, and TER119 antibodies. Following CD45-negative gating, cells were sorted 4 ways using a FACSAria as shown in (A); S1 and S2 were combined due to low cell number. S5, indicated in grey in Figure 1A, was not included in sorts due to low cell number and technical limitations. (B) Quantitative PCR was performed on sorted populations for indicated anti-apoptotic family members. Expression is represented as fold change relative to housekeeping gene (*Ubc*), n=12 embryos from 3 different litters. Mean plus or minus SEM; one-way ANOVA with α=0.05.

### EpoR-Cre-mediated *Mcl1*-deletion results in failure of erythropoiesis

To determine whether MCL-1 was important during early erythropoiesis, *Mcl1* conditional mice (hereafter referred to as *Mcl1*^F/F^) were bred to mice expressing the Cre-recombinase under control of the *EpoR* gene (EpoR-Cre)^23,30^. Expression of *EpoR* begins during erythroid burst-forming unit (BFU-E) stage, reaches highest expression at erythroid colony forming unit (CFU-E) stage, and then decreases^2,38^. EpoR-Cre mice induce over 90% recombination efficiency in the fetal liver, primarily in early erythroid progenitor cells^30,39^. Intercrosses did not produce any *Mcl1*^F/F^ EpoR-Cre+ mice indicating that loss of *Mcl1* in the erythroid lineage results in embryonic lethality (Table 1A). Notably, *Mcl1*^F/wt^ EpoR-Cre+ mice were found at expected frequency demonstrating that deletion of a single allele of *Mcl1* is tolerated.

**Table 1.**
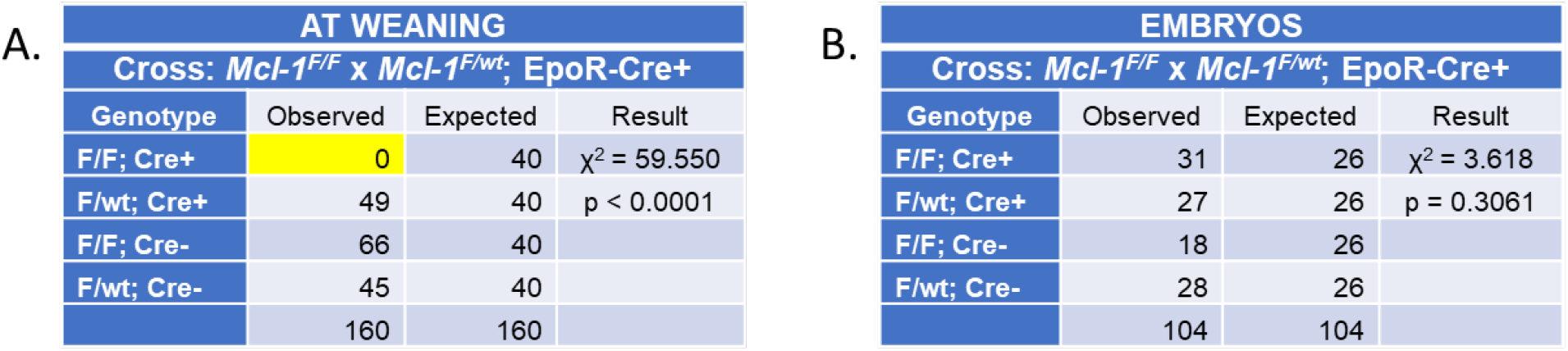
Genotypes of animals obtained from crosses between *Mcl1*^F^ and EpoR-Cre+ mice. (A) Result of crosses between *Mcl1*^F/F^ females and *Mcl1*^F/wt^; EpoR-Cre+ males with Chi-square analysis. Animals alive at weaning (average 3 weeks of age). Animals were genotyped upon weaning and Chi-square analyses were performed using Graphpad Prism version 6.02. As shown in the rightmost column of (A) no *Mcl1*^F/F^ EpoR-Cre+ animals survived to weaning of 280 total live births. These data indicate that erythroid-specific deletion of *Mcl1* is incompatible with life. (B) Embryos obtained from timed matings of *Mcl1*^F/F^ females with *Mcl1*^F/wt^; EpoR-Cre+ males (E12.5-13.5) were genotyped and Chi-square analyses were performed. *Mcl1*^F/F^ EpoR-Cre+ embryos are present in Mendelian ratios indicating that, at this time point, erythroid-specific loss of both alleles of *Mcl1* is tolerated.

| AT WEANING |  |  |  |
| --- | --- | --- | --- |
| Cross: *Mcl-1*<sup>F/F</sup> x *Mcl-1*<sup>F/wt</sup>; *EpoR-Cre*<sup>+</sup> |  |  |  |
| Genotype | Observed | Expected | Result |
| F/F; Cre+ | 0 | 40 | $\chi^2 = 59.550$ |
| F/wt; Cre+ | 49 | 40 | $p < 0.0001$ |
| F/F; Cre- | 66 | 40 |  |
| F/wt; Cre- | 45 | 40 |  |
|  | 160 | 160 |  |

B.
| EMBRYOS |  |  |  |
| --- | --- | --- | --- |
| Cross: *Mcl-1*<sup>F/F</sup> x *Mcl-1*<sup>F/wt</sup>; *EpoR-Cre*<sup>+</sup> |  |  |  |
| Genotype | Observed | Expected | Result |
| F/F; Cre+ | 31 | 26 | $\chi^2 = 3.618$ |
| F/wt; Cre+ | 27 | 26 | $p = 0.3061$ |
| F/F; Cre- | 18 | 26 |  |
| F/wt; Cre- | 28 | 26 |  |
|  | 104 | 104 |  |

To determine at which developmental stage the *Mcl1*^F/F^ EpoR-Cre+ mice were lost, embryos were obtained from timed pregnancies. In contrast to mice at weaning, at E13.5 we found that *Mcl1*^F/F^ EpoR-Cre+ embryos were observed at expected ratios (Table 1B). Phenotypically, the *Mcl1*^F/F^ EpoR-Cre+ embryos were smaller and paler than littermate controls when isolated at various time points (Figure 2A). Pathologically, fetal livers from erythroid *Mcl1*-deleted animals were smaller, exhibited ischemic necrosis, and lacked mature red blood cells (RBCs) in the sinusoids (Figure 2Bi&ii). Additionally, apoptotic cells were evident in fetal livers of *Mcl1*^F/F^ EpoR-Cre+ embryos when compared to littermate controls (Figure 2Bii, arrows). Circulating RBCs were reduced in *Mcl1*^F/F^ EpoR-Cre+ embryos and retained nuclei, unlike those observed in control *Mcl1*^F/wt^ EpoR-Cre+ embryos (Figure 2Biii). Similarly, cytospins from E12.5 fetal livers revealed that *Mcl1*^F/F^ EpoR-Cre+ erythroid cells retain their nuclei, unlike littermate control fetal livers, which exhibited mature, non-nucleated, erythroid cells (Figure 2C).

**Figure 2.**
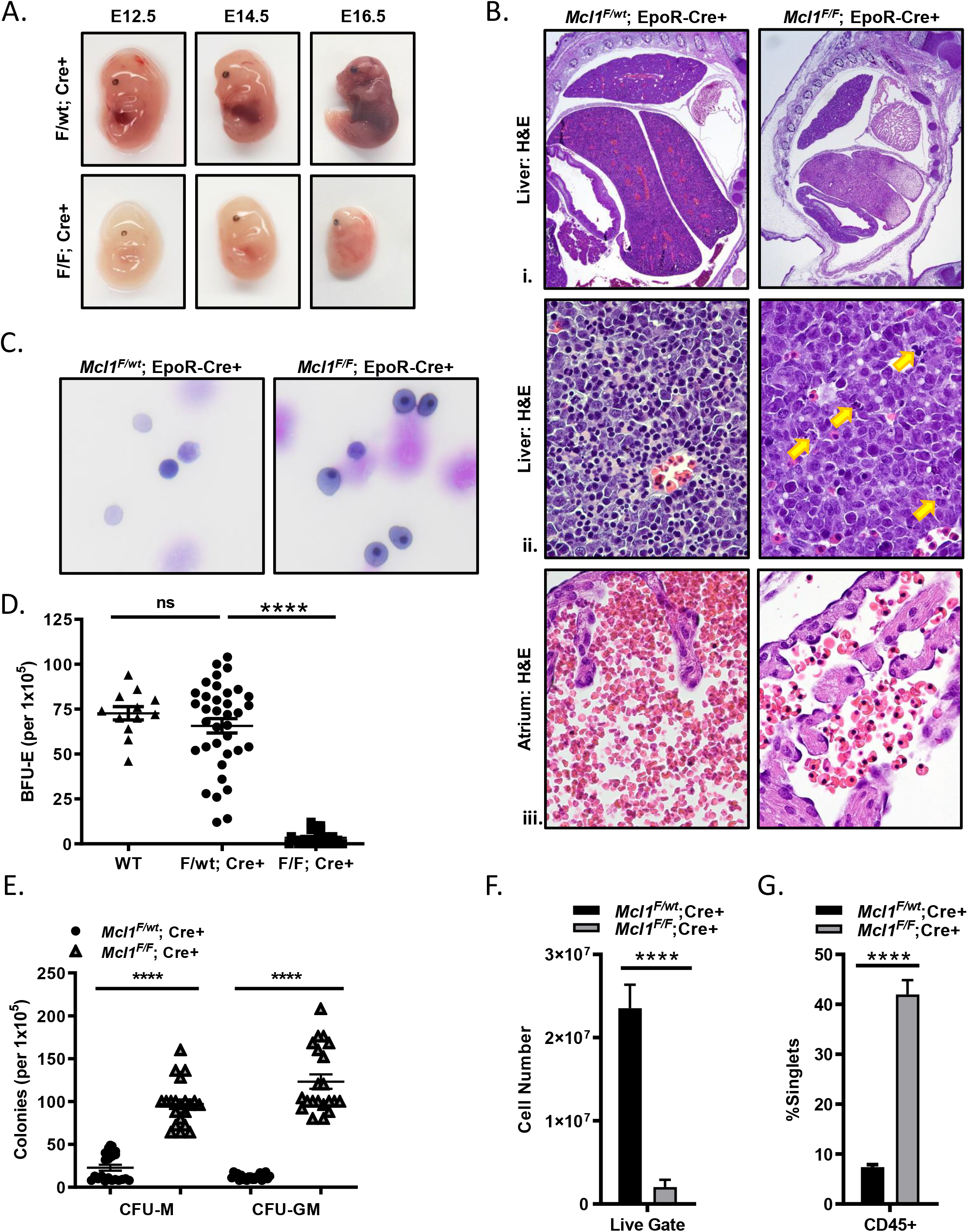
EpoR-Cre-mediated *Mcl1* deletion leads to lethal failure of blood development. (A) Photographs of freshly isolated representative *Mcl1^F/F^* EpoR-Cre+ and *Mcl1^F/wt^* EpoR-Cre+ embryos at days E12.5, E14.5, and E16.5. (B) Representative images of H&E-stained liver and heart sections from E12.5 embryos. (i) Liver imaged at 2x to indicate size and gross morphology; (ii) Liver imaged at 60x with yellow arrows indicating areas of apoptosis; (iii) Atrium imaged at 60x to indicate number and phenotype of RBCs. (C) Cytospin preparations from freshly-isolated fetal livers stained with DipQuick staining kit and imaged at 40x. (D, E) Methylcellulose colony-forming assays were performed using total FLC and medium as indicated; each dot represents a plate (n>12, mean plus or minus SEM, one-way ANOVA with α=0.05. **p < 0.1; and ****p < 0.001): (D) Enumeration of primitive erythroid progenitor cells (BFU-E) grown on MethoCult^™^ GF M3434 (contains Epo). (E) Enumeration of granulocyte-macrophage progenitor cells (CFU-M and CFU-GM) grown on MethoCult^™^ GF M3534 (without Epo). (F, G) Flow cytometric data from fetal liver cells (n=8-12 embryos from 4 different litters, mean plus or minus SEM, Student’s t Test. ****p < .001): (F) Enumeration of cells within the live gate. (G) Percent CD45-positive cells of singlets (Live gate > singlets > CD45 gate).

To examine the effects of EpoR-Cre-mediated loss of *Mcl1* on erythropoiesis, as well as on other hematopoietic lineages, E12.5 fetal liver cells were cultured in methylcellulose containing hematopoietic growth factors to assess colony formation^33,40^. Fetal liver cells from *Mcl1*^F/F^ EpoR-Cre+ embryos failed to produce erythroid lineage blast forming units (BFU-E), whereas mice lacking only one allele of *Mcl1* formed similar numbers of BFU-E to wild-type animals when cultured in the presence of Epo (Figure 2D).

The inability to give rise to BFU-E was not reflective of a general inability to produce hematopoietic colonies as *Mcl1*^F/F^ EpoR-Cre+ fetal liver cells gave rise to monocyte colony forming units (CFU-M) and mixed granulocyte/macrophage colony forming units (CFU-GM)(Figure 2E). These data indicate that fetal livers from *Mcl1*^F/F^ EpoR-Cre+ embryos are defective in definitive erythropoiesis, but capable of forming non-erythroid hematopoietic colonies.

### EpoR-Cre-mediated *Mcl1*-deletion does not perturb hematopoietic progenitors

While the EpoR-Cre expressing mouse has been commonly used to assess erythropoiesis, it has been reported that EpoR-Cre expression can occur even earlier in early hematopoietic progenitors^41^. Since MCL-1 is essential for promoting survival during early hematopoiesis22, we wanted to confirm that the loss of erythropoiesis was not due to earlier deletion in hematopoietic progenitors. The hematopoietic progenitors from E13.5 fetal livers were assessed by flow cytometry. Although the cellularity of *Mcl1*^F/F^ EpoR-Cre+ fetal livers was reduced in comparison to littermate controls, the percentages of hematopoietic progenitors was higher in the *Mcl1*^F/F^ EpoR-Cre+ embryos when compared to *Mcl1^F/wt^* EpoR-Cre+ littermate controls (Figure 2F&G and Sup. Fig. S1A and S1B). When progenitor numbers were compared directly, the total amount was similar or higher in the *Mcl1*^F/F^ EpoR-Cre+ fetal liver when compared to littermate controls (Sup. Fig. S1B and S1D). t-Stochastic Neighbor Embedding (t-SNE) mapping of hematopoietic progenitor populations from *Mcl1*^F/F^ EpoR-Cre+ fetal livers and littermate controls revealed that while some clusters appeared at different densities, no progenitor populations were missing in the *Mcl1*^F/F^ EpoR-Cre+ fetal livers (Sup. Fig. S1C). Further analysis confirmed the presence of downstream hematopoietic populations (Sup. Fig. S1E, F, G). These data indicate that the loss of definitive erythropoiesis in *Mcl1*^F/F^ EpoR-Cre+ embryos is not due to an earlier hematopoietic defect, but caused by *Mcl1*-deletion specifically during erythropoiesis.

### MCL-1 loss arrests development beyond early basophilic erythroblasts

To determine the stage during erythropoiesis when EpoR-Cre-mediated *Mcl1* deletion affects development, flow cytometry was used to evaluate fetal livers from timed pregnancies33. Analysis of control *Mcl1*^F/wt^ EpoR-Cre+ fetal livers revealed a stepwise progression from S0 (primitive progenitors) to S5 (late orthochromatophilic erythroblasts and reticulocytes) over 4 days of fetal liver development from E11.5-E15.5 (Figure 3A). Conversely, in *Mcl1*^F/F^ EpoR-Cre+ fetal livers, the preponderance of cells were restricted to the S0 and S1 populations. Few *Mcl1*^F/F^ EpoR-Cre+ fetal liver cells progressed beyond the S1 (proerythroblast) stage even when harvested as late as E15.5 (Figure 3A). Furthermore, unlike *Mcl1*^F/wt^ EpoR-Cre+, which exhibited increased fetal liver cellularity as development progressed through the S3 (late basophilic erythroblasts) and S4 (chromatophilic and early orthochromatophilic erythroblasts) stages, numbers of *Mcl1*^F/F^ EpoR-Cre+ fetal liver erythroid progenitors were reduced at all time points analyzed (Figure 3B). The reduction in cellularity was in part due to increased apoptotic (Annexin-V^+^) erythroid progenitors in the *Mcl1*^F/F^ EpoR-Cre+ fetal livers, while deletion of only a single allele of *Mcl1* did not increase apoptotic cells (Figure 3C).

**Figure 3.**
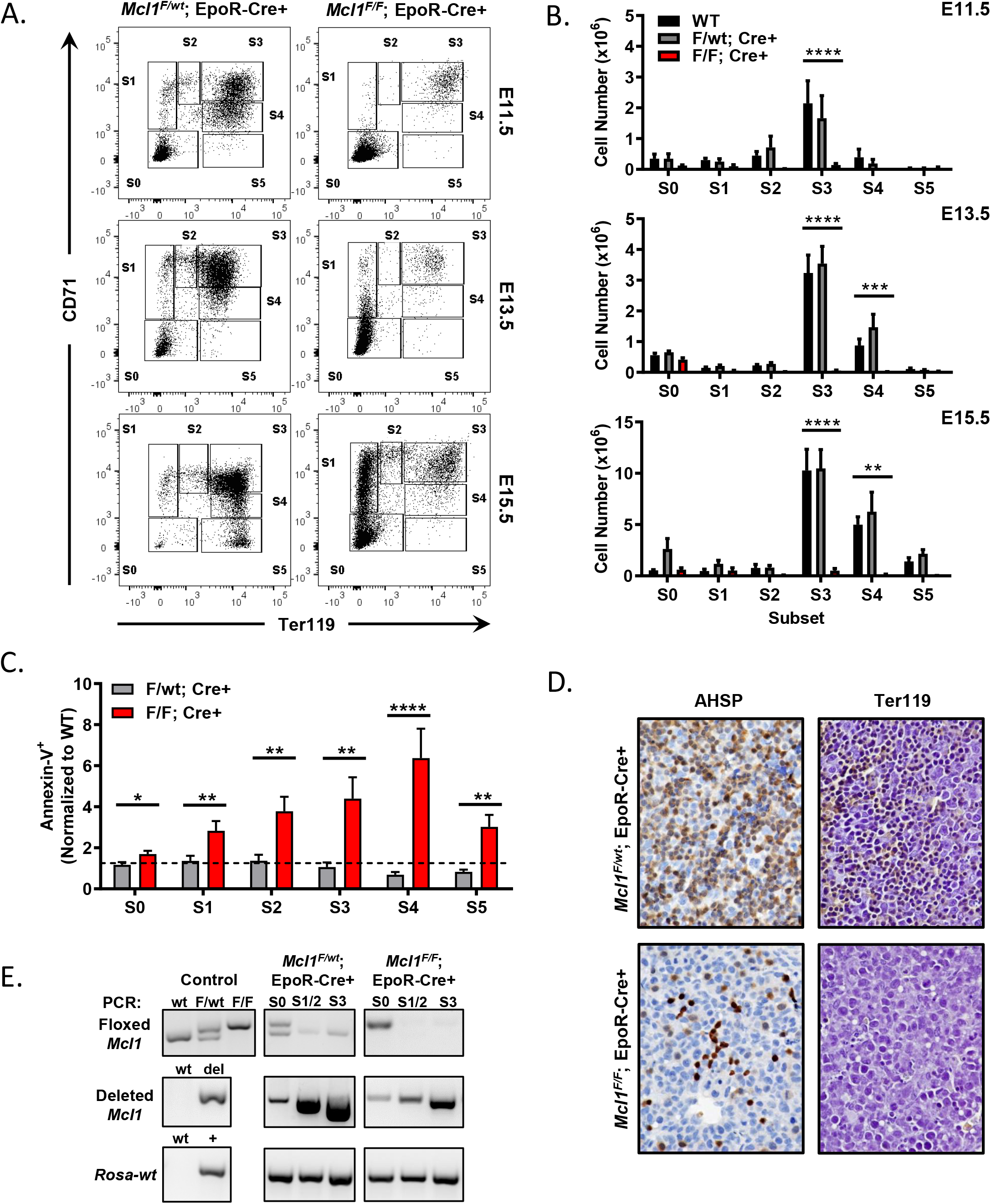
EpoR-Cre-mediated deletion of *Mcl1* leads to blockade early in erythrocyte development. (A-C) Mouse fetal liver cells were isolated from *Mcl1^F/F^* EpoR-Cre+ or *Mcl1^F/wt^* EpoR-Cre+ embryos at the indicated day, labeled with CD45, CD71, TER119, and Annexin-V antibodies, and analyzed by flow cytometry. (A) CD45-negative cells representing E11.5 (top row), E13.5 (middle), and E15.5 (bottom) embryos are shown. (B) Absolute cell numbers present in each differentiation stage as represented in (A); E11.5 (top row), E13.5 (middle), and E15.5 (bottom) are shown (n=at least 5 biological replicates with 4 embryos each, mean plus or minus SEM, One-way ANOVA with α=0.05. Not indicated, not significant; **p<0.1; ***p<0.01; ****p < 0.001). (C) Annexin-V-positive (dead) cells in E13.5 *Mcl1^F/F^* EpoR-Cre+ (red) or *Mcl1^F/wt^* EpoR-Cre+ (grey) fetal livers. Graph represents percent relative to wildtype control (n>13 embryos from 5 different litters, Student’s t Test. *p<0.5; **p<0.01; ****p <0.001). (D) Liver sections from representative embryos subjected to immunohistochemical (IHC) staining for alpha hemoglobin stabilizing protein (AHSP, left) and a glycophorin A-associated protein (TER119, right). Images are 60x, see Methods for further details. (E) DNA from fetal liver cells isolated from *Mcl1^F/F^* EpoR-Cre+ or *Mcl1^F/wt^* EpoR-Cre+ embryos were analyzed by PCR; “Control” indicates PCR control and represents expected banding pattern.

E12.5 *Mcl1^F/F^* EpoR-Cre+ fetal livers had low TER119 expression when compared to littermate control *Mcl1*^F/wt^ EpoR-Cre+ fetal liver sections (Figure 3D); TER119 is an erythroid-specific antigen whose expression starts on early pro-erythroblasts^42^. Furthermore, *Mcl1*^F/F^ EpoR-Cre+ fetal livers also exhibited fewer cells expressing alpha hemoglobin stabilizing protein (AHSP, Figure 3D), a molecular chaperone which prevents harmful aggregation and precipitation of free alpha-globin from occurring during normal erythropoiesis^43^. In normal erythropoiesis, AHSP expression is first detected in the proerythroblast stage, peaks at the orthochromatic stages, and declines to low levels in polychromatophilic reticulocytes^44,45^. Low expression of TER119 and aberrant expression of AHSP together indicate a severe defect in erythroid differentiation^46^.

To assess the stage at which EpoR-Cre was inducing recombination, we performed PCR on DNA from flow-sorted erythroid progenitors to detect the *Mcl1* and deleted *Mcl1* alleles. In both *Mcl1*^F/wt^ EpoR-Cre+ and *Mcl1*^F/F^ EpoR-Cre+ fetal livers there was some evidence of recombined *Mcl1* as early as the S0 stage of erythropoiesis, but by the S1/2 stages the un-recombined floxed *Mcl1* genomic allele was absent indicating efficient deletion (Figure 3E). Taken together, these data indicate that EpoR-Cre activity initiates during the S0 stage, but was most efficient at mediating recombination of *Mcl1* at the S1/2 stages of differentiation.

### MCL-1 promotes early erythropoiesis but is dispensable during later stages

*Mcl1*^F/F^ EpoR-Cre+ fetal livers indicate that MCL-1 is required during the early stages of erythroid differentiation; however, these experiments are unable to assess whether MCL-1 plays similar critical roles later during erythropoiesis. To control the timing of *Mcl1* deletion, we utilized fetal livers from *Mcl1^F/F^* Rosa-ERCreT2 mice in which 4-OH-tamoxifen (4-OHT) can be added to the culture media to induce Cre-mediated recombination at specified times during culture. TER119-depleted E12.5 fetal liver cells isolated from *Mcl1*^F/F^ Rosa-ERCreT2+ or *Mcl1*^F/wt^ Rosa-ERCreT2+ littermates were incubated for 18 hours in “stem cell media,” which contains SCF, IL-6, and FLT-3L to maintain progenitors, after which they were washed and re-plated in “differentiation medium” containing erythropoietin to promote erythroid differentiation^33,34^.

To induce *Mcl1*-deletion at the earliest stages of erythropoiesis, 4-OHT was added immediately with “stem cell media” to induce recombination early during culture (indicated as “time=0”). In contrast, to assess MCL-1’s function at later stages of erythropoiesis, deletion was induced by adding 4-OHT after 24 hours in culture, when the cells have been switched into erythropoietin-containing media and were already in the process of erythroid differentiation (indicated as “time=24”). PCR of bulk cultures showed deletion of the floxed *Mcl1* allele at both time points (Figure 4A), while immunoblotting showed a clear decrease in MCL-1 protein in *Mcl1*^F/F^ Rosa-ERCreT2+ cells following 4-OHT treatment (Figure 4B). Both untreated *Mcl1*^F/F^ Rosa-ERCreT2+ or *Mcl1*^F/wt^ Rosa-ERCreT2+ littermate control fetal liver cells were able to efficiently differentiate as assessed by CD71 and TER119 staining (Figures 4C&E). Early addition of 4-OHT to *Mcl1*^F/F^ Rosa-ERCreT2+ fetal liver cells (time=0) resulted in >50% of cells that failed to progress beyond the S0 stage (Figures 4C&F); however, later addition of 4-OHT (time=24) did not alter erythropoiesis relative to the untreated condition (Figures 4C&G). Cytospins from these cultures revealed that *Mcl1*^F/wt^ Rosa-ERCreT2+ fetal liver cells were able to differentiate regardless of when 4-OHT is added, indicating that deletion of a single *Mcl1* allele was not deleterious (Figure 4D). In contrast, *Mcl1^F/F^* Rosa-ERCreT2+ treated with 4-OHT at t=0 generated mostly debris and progenitor cells, while the same cells treated with 4-OHT at t=24 differentiated normally and generated enucleated erythrocytes (Figure 4D & Sup. Figure S2). Annexin-V staining indicated that >80% of *Mcl1^F/F^* Rosa-ERCreT2+ fetal liver cells treated with 4-OHT at time=0 were apoptotic, suggesting that the role for MCL-1 during the early stages of erythropoiesis is to promote survival (Figure 4H).

**Figure 4.**
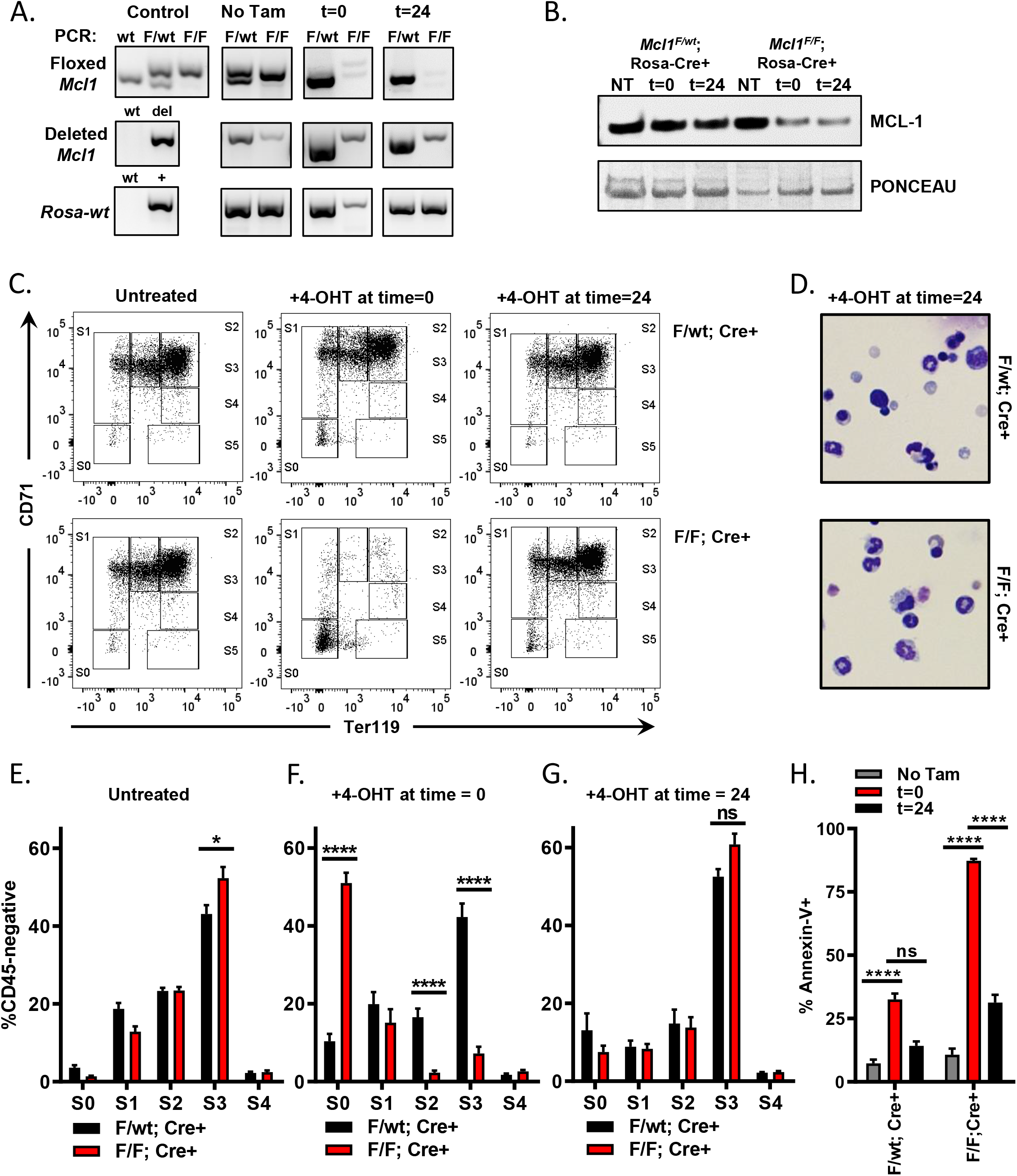
*Ex vivo* culture system allows temporally controlled deletion of *Mcl1* from erythroid cultures. Fetal liver cells were isolated from E12.5 *Mcl1^F/F^* Rosa-ERCreT2+ or *Mcl1^F/wt^* Rosa-ERCreT2+ embryos; TER119-negative progenitor cells were purified using MACS microbeads then cultured on fibronectin-coated plates. To induce *Mcl1* deletion, FLC were treated with 4-OH-Tamoxifen (4-OHT) during the first 18 hours/overnight of culture (t=0) or overnight the second day of culture (t=24). Controls did not receive 4-OHT (“No Tam”). (A) Cells were cultured as indicated, then harvested and analyzed by PCR. “Control” indicates PCR control and represents expected banding pattern while “No Tam” represents experimental control and represents non-deleted cells. (B) Immunoblot of MCL-1 levels in *in vitro* differentiated cells treated with 4-OHT as indicated (NT = No tamoxifen). Ponceau-S staining was used to indicate loading. (C) Representative flow cytometric analysis of CD45-negative *in vitro* differentiated FLC. (D) Cytospin preparations from *Mcl1^F/wt^* Rosa-ERCreT2+ or *Mcl1^F/F^* Rosa-ERCreT2+ FLC treated with 4-OHT overnight the second day of culture stained with DipQuick staining kit and imaged at 40x. (E-G): Quantification of *Mcl1^F/wt^* Rosa-ERCreT2+ (black) and *Mcl1^F/F^* Rosa-ERCreT2+ (red) *ex vivo* differentiated fetal liver cells by flow cytometry (n>12 embryos from 3 different litters, mean plus or minus SEM, One-way ANOVA with α=0.05. ns or not indicated, not significant; *p<0.05; ****p < 0.001): (E) Differentiation stages in untreated control cells. (F) Differentiation stages in cells treated with 4-OHT during the first 18 hours of culture. (G) Differentiation stages in cells treated with 4-OHT overnight the second day of culture. (H) Cells were differentiated as indicated, then harvested and labeled with CD45, CD71, TER119, and Annexin-V antibodies. Graph indicates percent Annexin-V-negative (alive) cells (n>6 embryos from 2 biological replicates, mean plus or minus SEM, One-way ANOVA with α=0.05. ns or not indicated, not significant; *p<0.05; ****p < 0.001).

### Removal of apoptosis rescues *Mcl1* deficiency in a transplant model

In order to investigate if *Mcl1* exerts its function by inhibiting apoptosis, we sought to co-delete *Mcl1* along with the essential pro-apoptotic effector molecules *Bax* and *Bak.* Since cells lacking *Bax* and *Bak* are unable to undergo canonical apoptosis^47^, we hypothesized that removal of *Mcl1* on this background should rescue erythropoiesis if MCL-1’s critical function is to restrain apoptosis. To this aim, lethally irradiated CD45.1^+^ recipient animals were transplanted with whole BM from the donor groups shown in Figure 5A. After 4 weeks of engraftment, tamoxifen was administered to the recipient animals to induce deletion of *Mcl1* and/or *Bax* conditional alleles on a *Bak*-deficient background. Complete blood counts (CBC) and flow cytometry were conducted every two weeks after tamoxifen administration showed a decrease in neutrophils and a concomitant increase in lymphocytes following tamoxifen treatment, but these trends are consistent across all cohorts (Sup. Figure S3B, D, F&H).

**Figure 5.**
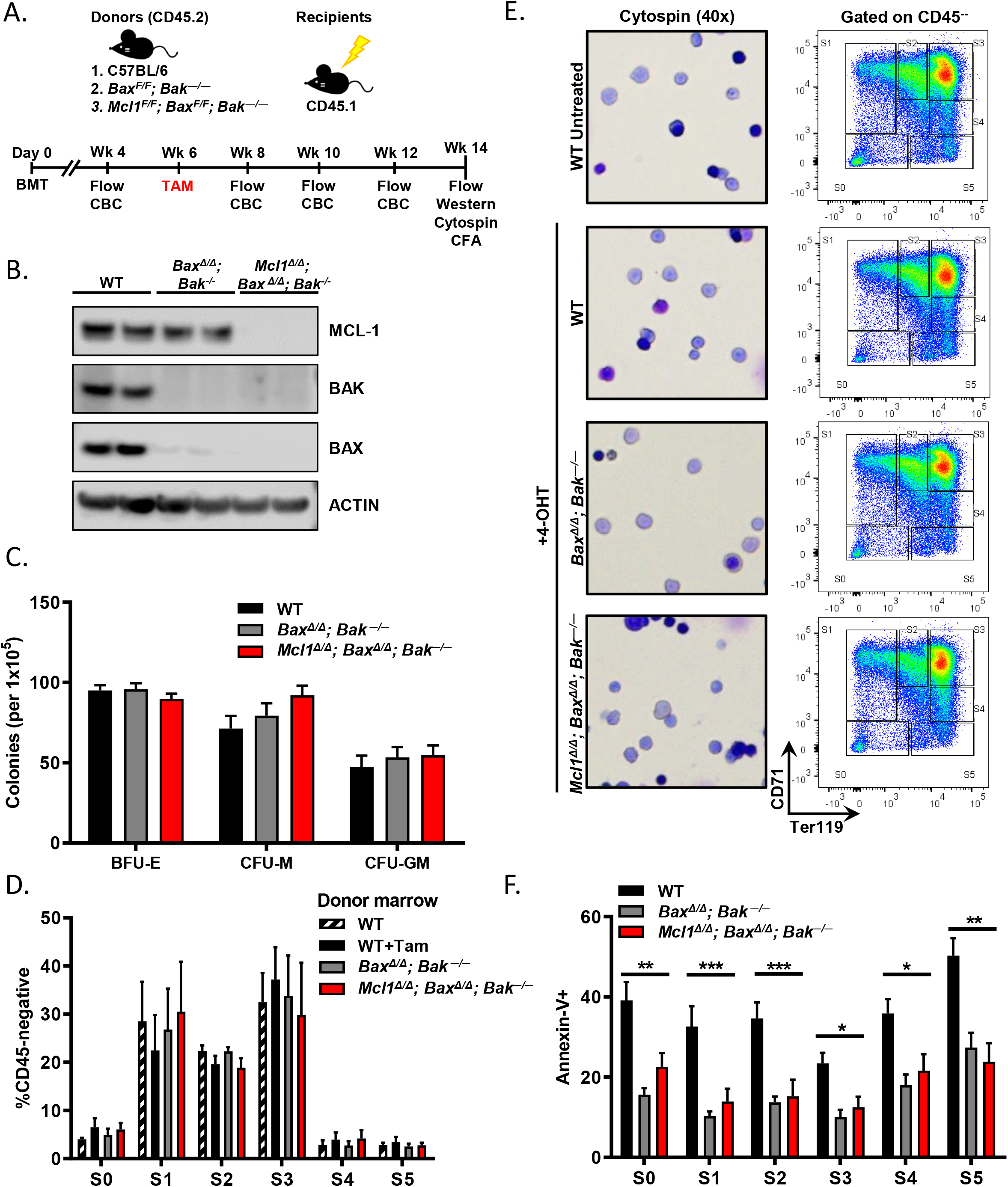
Removal of apoptosis rescues erythropoiesis defect caused by deletion of *Mcl1*. (A) Schematic of transplant studies indicating lethally irradiated CD45.1+ recipients and three separate groups of CD45.2+ donors: C57BL/6 wildtype, *Bax^F/F^Bak^−/−^Rosa*-ERCreT2, and *Mcl1^F/F^ Bax^F/F^ Bak^−/−^* Rosa-ERCreT2 (2 biological replicates with 5-6 recipients per donor group, see Methods). (B) Immunoblot of recipient spleens (2 lanes each) following bone marrow (BM) transplant and tamoxifen-mediated deletion of floxed alleles. (C, D, F) Recipient BM was harvested and differentiated into erythroblasts on fibronectin-coated plates as in Figure 4. (C) *Ex vivo* differentiated recipient BM cells were labeled with CD45, CD71, and TER119 then analyzed by flow cytometry. CD45-negative fraction is shown. (D) Representative cytospin preparations from *ex vivo* differentiated recipient BM (left), and flow cytometry histograms (gated on CD45-negative fraction). (E) Methylcellulose colony-forming assays were performed using recipient total BM; primitive erythroid progenitor cells (BFU-E) were grown on MethoCult^™^ GF M3434 (contains Epo) while granulocyte-macrophage progenitor cells (CFU-M and CFU-GM) were grown on MethoCult^™^ GF M3534 (without Epo). (F) *Ex vivo* differentiated BM was labeled with CD45, CD71, TER119, and Annexin-V antibodies and analyzed by developmental stage (n=5-6 recipient animals per donor group, 2 biological replicates each. Mean plus or minus SEM, Two-way ANOVA with α=0.05. *p<0.05; **p<0.1; ***p < 0.01).

Six weeks after tamoxifen administration, hematopoietic tissues were examined. As expected, mice receiving wild-type marrow showed robust expression of all three proteins (Figure 5B). Mice receiving *Bax^F/F^Bak*^−/−^Rosa-ERCreT2 marrow expressed MCL-1, and those receiving *Mcl1*^F/F^*Bax^F/F^Bak*^−/−^Rosa-ERCreT2 marrow were devoid of BAX,BAK, and MCL-1, indicating effective transplant and tamoxifen-mediated deletion of the expected floxed alleles (Figure 5B).

To assess erythropoiesis, we subjected harvested BM from the tamoxifen-treated recipient mice to *ex vivo* differentiation culture systems (as described for fetal liver). When BM from recipients was cultured on methylcellulose for colony forming assays, wildtype and *Bax*^Δ*/*Δ^*Bak*^−/−^Rosa-ERCreT2 BM efficiently differentiated into BFU-E, CFU-M, and CFU-GM, demonstrating that *Bax*- and *Bak*-deficiency does not compromise colony formation (Figure 5C). Similarly, *Mcl1*^Δ/Δ^*Bax*^Δ/Δ^*Bak*^−/−^Rosa-ERCreT2 cells also generated BFU-E when cultured in Epo-containing methylcellulose and differentiated into CFU-M and CFU-GM (Figure 5C). These data indicate that co-deletion of *Bax* and *Bak* rescued erythroid differentiation in the absence of *Mcl1* without any deleterious effect on myeloid differentiation (Sup. Figure S3). Furthermore, recipient marrow was isolated, lineage-depleted, and cultured on fibronectin-coated plates in Epo-containing media. Donor marrow samples from each genotype were able to efficiently differentiate into erythroid cells as indicated by loss of CD45 and expression of erythroid markers (Figures 5D&E); there was no significant difference among groups. Cytospins confirmed the presence of mature, enucleated erythrocytes in cultures derived from all tested genetic backgrounds (Figure 5E). Finally, the *ex vivo* differentiated BM was stained with Annexin-V; as expected, *Bax*^Δ/Δ^*Bak*^−/−^Rosa-ERCreT2 BM exhibited a lower frequency of apoptotic cells relative to WT controls (Figure 5F). However, *Mcl1*^Δ/Δ^*Bax*^Δ/Δ^*Bak*^−/−^Rosa-ERCreT2 cells from the erythroid cultures displayed significantly diminished apoptosis at all stages of erythroid development when compared to their wild-type counterparts (Figure 5F). These data indicate that hematopoiesis can occur normally despite loss of MCL-1 if the canonical apoptotic mediators BAX and BAK are ablated; taken together, these data suggest that MCL-1 functions to protect early stage erythrocytes from apoptosis.

### Ectopic *BCL2* expression rescues *Mcl1*-deficiency in an *ex vivo* differentiation model

Removal of both apoptotic effectors can compensate for loss of MCL-1, suggesting that MCL-1’s main contribution during erythropoiesis is promoting survival. However, it is still unclear whether MCL-1 is essential during early erythropoiesis simply because it is expressed early prior to the induction of other anti-apoptotic regulators, like BCL-xL (Figure 1B). Alternatively, MCL-1 may play a critical function in promoting survival that cannot be compensated for by other anti-apoptotic molecules. To differentiate between these hypotheses, we tested if the ectopic expression of another anti-apoptotic BCL-2 family member, human *BCL2*, could compensate for the deletion of *Mcl1* during *ex vivo* erythroid culture. TER119-depleted fetal liver cells from E12.5 *Mcl1*^F/F^ Rosa-ERCreT2+ or *Mcl1*^F/wt^ Rosa-ERCreT2+ littermate controls were infected with lentivirus containing human cDNAs encoding *BCL2, MCL1*, or negative control *EGFP* for 16 hours in the presence of SCF, IL-6, and Flt-3L to maintain stemness. During this time, 4-OHT was added to induce deletion of endogenous *Mcl1*. After 16 hours, the media was replaced with Epo-containing media to induce the erythroid differentiation program.

Since the lentiviral vectors co-expressed ZsGreen, flow cytometry was performed within 24 hours after transduction to assess efficiency of transduction (Figure 6A&B and Sup. Figure S4C). Successful transduction was confirmed by amplification of the human cDNAs encoded for by each lentivirus from cultured erythrocytes as well as the efficiency of *Mcl1* locus deletion (Figure 6C). Immunoblot was used to confirm expression of full-length human proteins from transduction using the same viruses in the murine 3T3 cell line (Sup. Figure S4A). Non-deleted fetal liver cells generated erythroid cells regardless of whether they expressed *BCL2* or *Mcl1*, indicating that ectopic expression did not hamper erythroid differentiation (Figure 6D).

**Figure 6.**
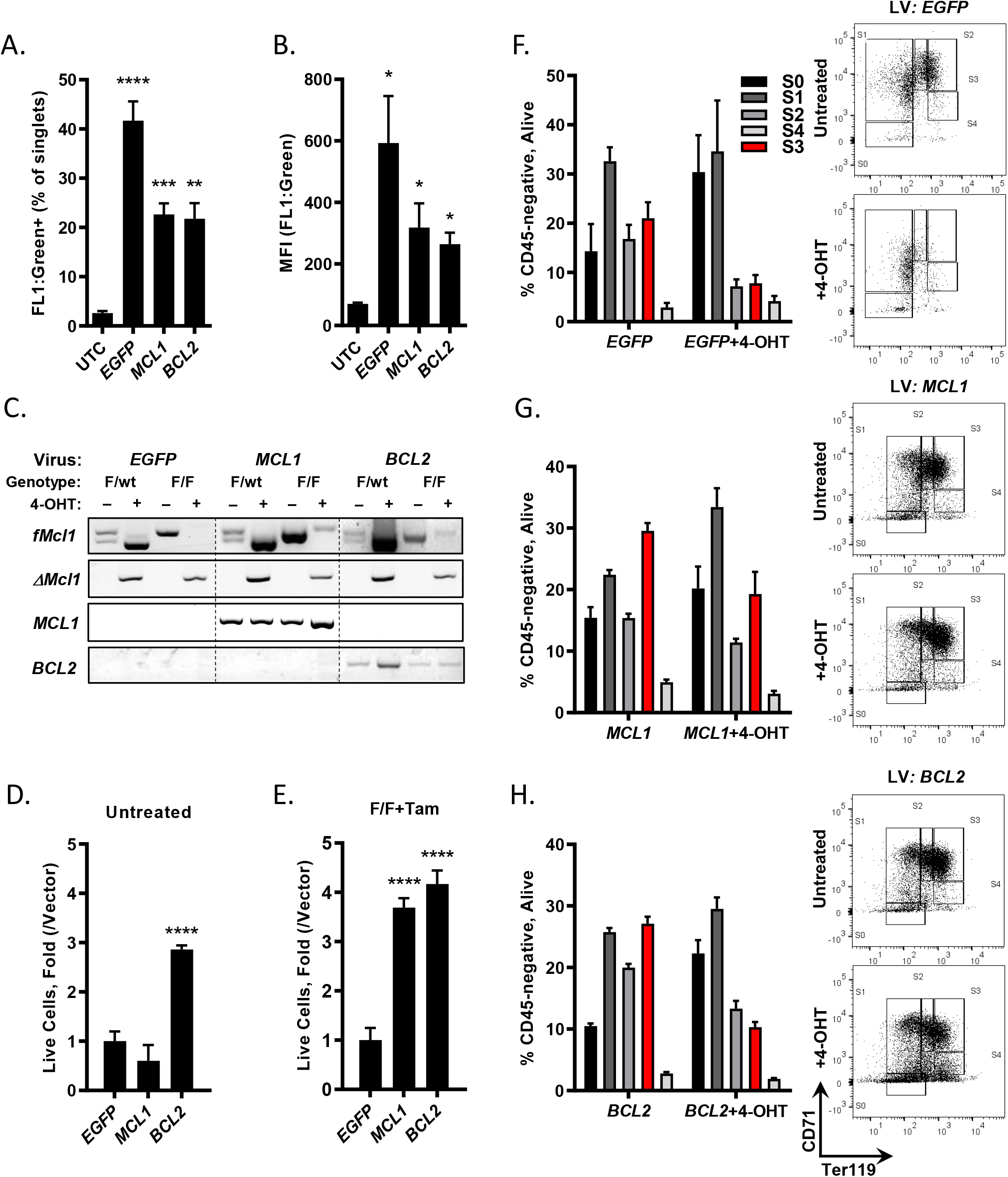
Lentivirus-mediated expression of human BCL-2 can rescue loss of MCL-1 in early erythropoiesis. (A-H) TER119-depleted fetal liver cells from E12.5 *Mcl1*^F/F^ Rosa-ERCreT2+ or Mcl1^F/WT^ Rosa-ERCreT2+ littermate controls were infected with zsGreen-tagged lentivirus encoding human cDNAs *(BCL2, MCL1)* or *EGFP* for 16 hours. Cultures were pulsed with 4-OHT for the following 16 hours, then media was changed to Epo-containing media to support erythroid differentiation. (A-B) Flow cytometric analysis of viral vector expression in fetal liver cells; UTC, untransfected control cells. (A) Percent zsGreen positive cells in the singlet gate. (B) Mean fluorescence intensity (MFI) of zsGreen signal. (n=at least 3 biological replicates with 3 embryos each, mean plus or minus SEM, One-way ANOVA with α=0.05. *p<0.5; **p<0.01; ***p<0.01; ****p <0.001). (C) Fetal liver cells were infected with virus indicated, pulsed with 4-OHT as indicated, then harvested and analyzed by PCR. First row is floxed *Mcl1*; second row is deleted *Mcl1* where presence of band indicates deletion. Third row is the human lentivirus insert encoding *MCL1*; fourth row is the human lentivirus insert encoding *BCL2*. (D-E) Counts of lentivirus-infected cells following erythroid differentiation protocol. Cells were left untreated (D) or pulsed with 4-OHT (E), then harvested and counted. Counts are represented as fold change relative to empty vector. (n=at least 3 biological replicates with 3 embryos each, mean plus or minus SEM, One-way ANOVA with α=0.05. ns or not indicated, not significant; ****p <0.001). (F-H) Cells infected with the indicated viruses were differentiated, harvested, and analyzed by flow cytometry. CD45-negative cells falling into the live gate were analyzed by expression of CD71 and TER119. (E) Representative histograms (right) of empty vector-expressing cells left untreated (top) or pulsed with 4-OHT (bottom); graph shows cumulative differentiation stage data. (F) *MCL1*-expressing and (G) *BCL2*-expressing cells. (n=at least 3 biological replicates with 3 embryos each, mean plus or minus SEM.)

To test whether other anti-apoptotic molecules could compensate for *Mcl1*-deletion, experiments were performed with 4-OHT to assess if erythroid progenitors expressing virally encoded cDNAs were able to undergo differentiation (Figure 6F-H). EGFP expression alone was not able to promote erythroid differentiation when *Mcl1* was deleted; the cultures were stuck in S1, comparable to *Mcl1^F/F^* EpoR-Cre fetal liver cells (Figure 6E&F). In contrast, expression of ectopic human *MCL1* efficiently rescued erythroid differentiation in 4-OHT treated cultures indicating that human MCL-1 can compensate for deletion of endogenous *Mcl1* (Figure 6E&G). Ectopic expression of *BCL2* supported the expansion and differentiation of erythroid cells treated with 4-OHT (Figure 6E&H). Cytospins of reconstituted fetal liver cells treated with 4-OHT indicated that while control (EGFP) expression did not rescue erythrocyte development, expression of human BCL-2 or MCL-1 resulted in efficient erythroid differentiation like that observed in cells without 4-OHT (Sup. Figure S4B). These data demonstrate that human *BCL2* can functionally compensate for the *Mcl1*-deletion during erythroid differentiation, indicating that MCL-1 does not have a unique functional role in erythroid differentiation that cannot be provided by human BCL-2.

## Discussion

BCL-xL maintains the survival of mature erythrocytes and has been linked to signal transduction downstream of Epo^6,10,48^. Despite this knowledge, it was unclear which pro-survival factors were required to inhibit apoptosis during early erythroid development^49^. We found that *Mcl1-deletion* during the earliest stages of erythropoiesis resulted in a failure to develop beyond the S1/S2 stages, caused the apoptotic death of developing erythrocytes, and led to mid-gestational embryonic lethality. Phenotypically, *Mcl1* ablation in erythroid progenitors looks strikingly like *Epo*- or *EpoR-deficient* mice^2^. In contrast, deletion of *Bcl2l1* (encodes BCL-xL) leads to defective erythroid differentiation at a later stage of development^10-11,48^. These data indicate that MCL-1 plays a critical survival role during the early stages of erythropoiesis, when Epo signaling provides the survival cue.

Our findings indicate that developing erythrocytes switch their dependency on pro-survival BCL-2 family members. During the earliest stages of erythroid differentiation, endogenous MCL-1 provides an essential anti-apoptotic function to promote cell survival, but during the later stages of erythroid differentiation, MCL-1 becomes dispensable as BCL-xL expression increases. During the S1/S2 stages of erythroid differentiation, MCL-1 mRNA expression is highest among the family, but with subsequent differentiation, BCL-xL mRNA expression increases in agreement with previous observations^5,6^. This modulation of expression correlates well with the switch in functional dependency that we observed in an *in vitro* differentiation assay.

It is still unclear why developing erythrocytes switch between MCL-1 and BCL-xL during erythropoiesis. One possible explanation is that the molecules have different functional roles beyond blocking cell death. Both MCL-1 and BCL-xL function not only to prevent cell death but can facilitate mitochondrial function^50-52^. While this may be the case in other cell lineages, our data demonstrate that *Bax* and *Bak* co-deletion can overcome the requirement for MCL-1 during early erythropoiesis. Similarly, genetic loss of both *Bax* and *Bak* rescued *Bcl2l1*-deletion during erythroid development^53^. Together, these data suggest that MCL-1 and BCL-xL function primarily to prevent cell death during erythropoiesis and any other functional contributions are dispensable.

MCL-1 is distinct from other BCL-2 family members, possessing a predominately unstructured amino-terminus containing regulatory regions54. It was possible that this region contributed to erythroid differentiation; however, ectopic expression of human BCL-2, which lacks the unstructured domain found in MCL-1, compensated for MCL-1 during early erythropoiesis. These data further support the notion that MCL-1’s primary function in erythroid cultures is to prevent apoptosis. While human BCL-2 expression in differentiation cultures can overcome the endogenous requirement for MCL-1, it remains to be seen whether replacement of *BCL2* for *Mcl1* will trigger other developmental issues in mammals. Furthermore, these data do not exclude the possibility that MCL-1 may possess additional unidentified functions in other cell types.

The finding that MCL-1 is required to protect early erythroid precursors adds to the variety of hematopoietic cell lineages that depend upon MCL-149. While it is still unclear why MCL-1 plays such a broad role, one potential reason is that MCL-1 undergoes rapid, proteasome-mediated degradation^36-55,56^. This triggers robust, basal turnover in most cell types; however, post-translational modifications including phosphorylation and ubiquitinylation can enhance its stability or potentiate its turnover^56-58^. MCL-1’s rapid turnover and ability for signal transduction pathways to regulate expression levels makes it an ideal regulator of survival, poised to rapidly respond to extracellular signaling^54^. Accordingly, MCL-1 maintains the survival of erythroid progenitors during early erythropoiesis and likely responds to extracellular cues, such as Epo signaling. Future research should ascertain whether Epo signaling directly regulates MCL-1 expression and whether this occurs via transcriptional or post-transcriptional mechanisms.

## Acknowledgments

We thank the members of the Opferman laboratory and M. Kundu (SJCRH) for helpful discussions and for providing EpoR-Cre mice. We also thank the St. Jude Children’s Research Hospital Animal Resource Center, Veterinary Pathology Core, and Flow Cytometry and Cell Sorting Shared Resource for their support of this project. This project was supported by the National Institute of Health R01HL123543 and R01CA201069 (J.T.O); a Cancer Center Support Grant P30CA021765; and the American Lebanese Syrian Associated Charities (ALSAC) of St. Jude.

## Authorship

J.T.O. P.N. and M.E.T. conceived the study, J.T.O. and M.E.T. designed the experiments, and wrote the manuscript. M.E.T., E.K., K.H.S., and B.J.K. performed experiments, analyzed data, and prepared figures. R.A.A. and J.D.L. provided flow cytometry expertise and analysis. P.V. performed pathology imaging and analysis. J.T.O. supervised the project.

## Supplementary Figure Legends

**Supplemental Figure S1.**
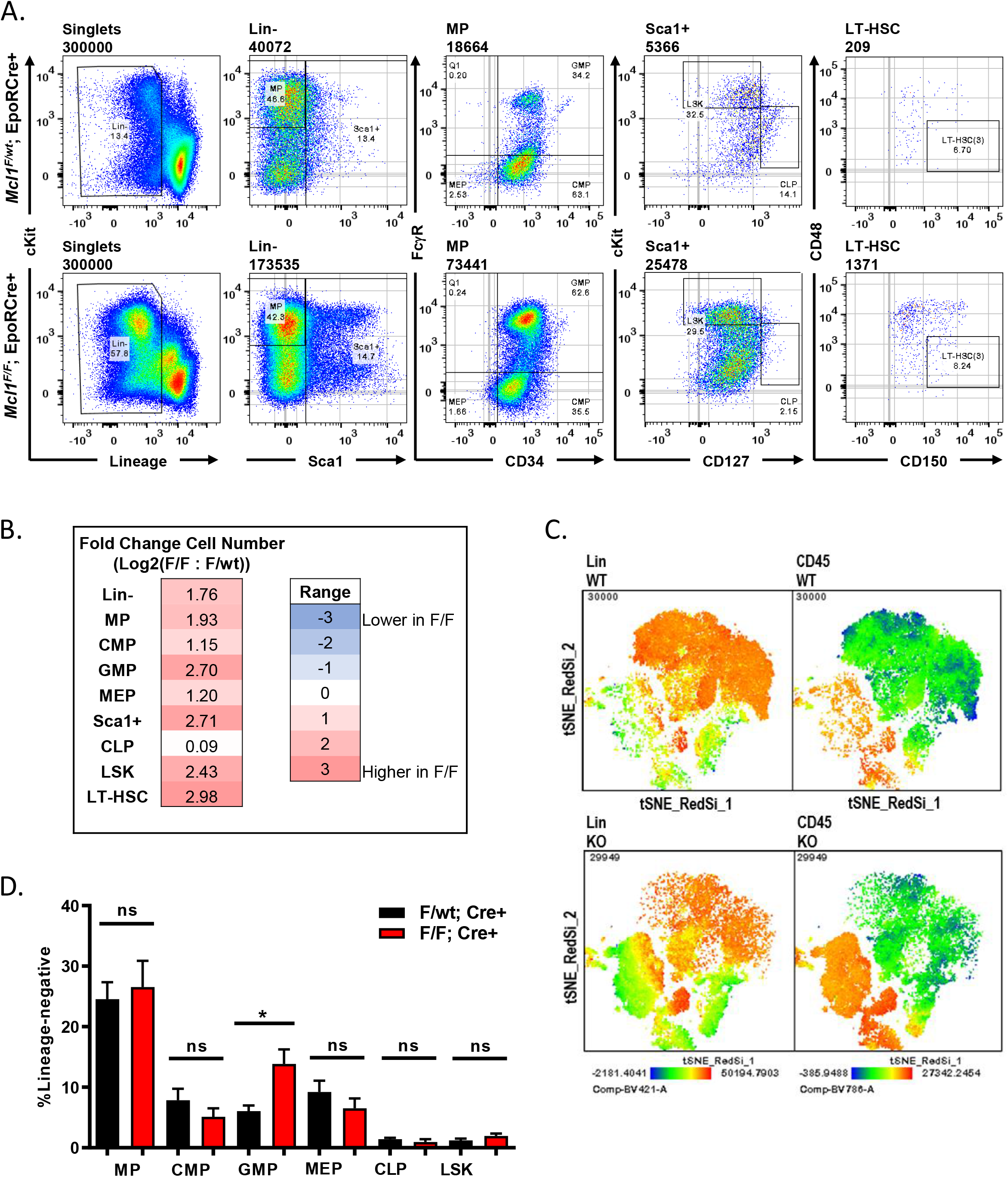
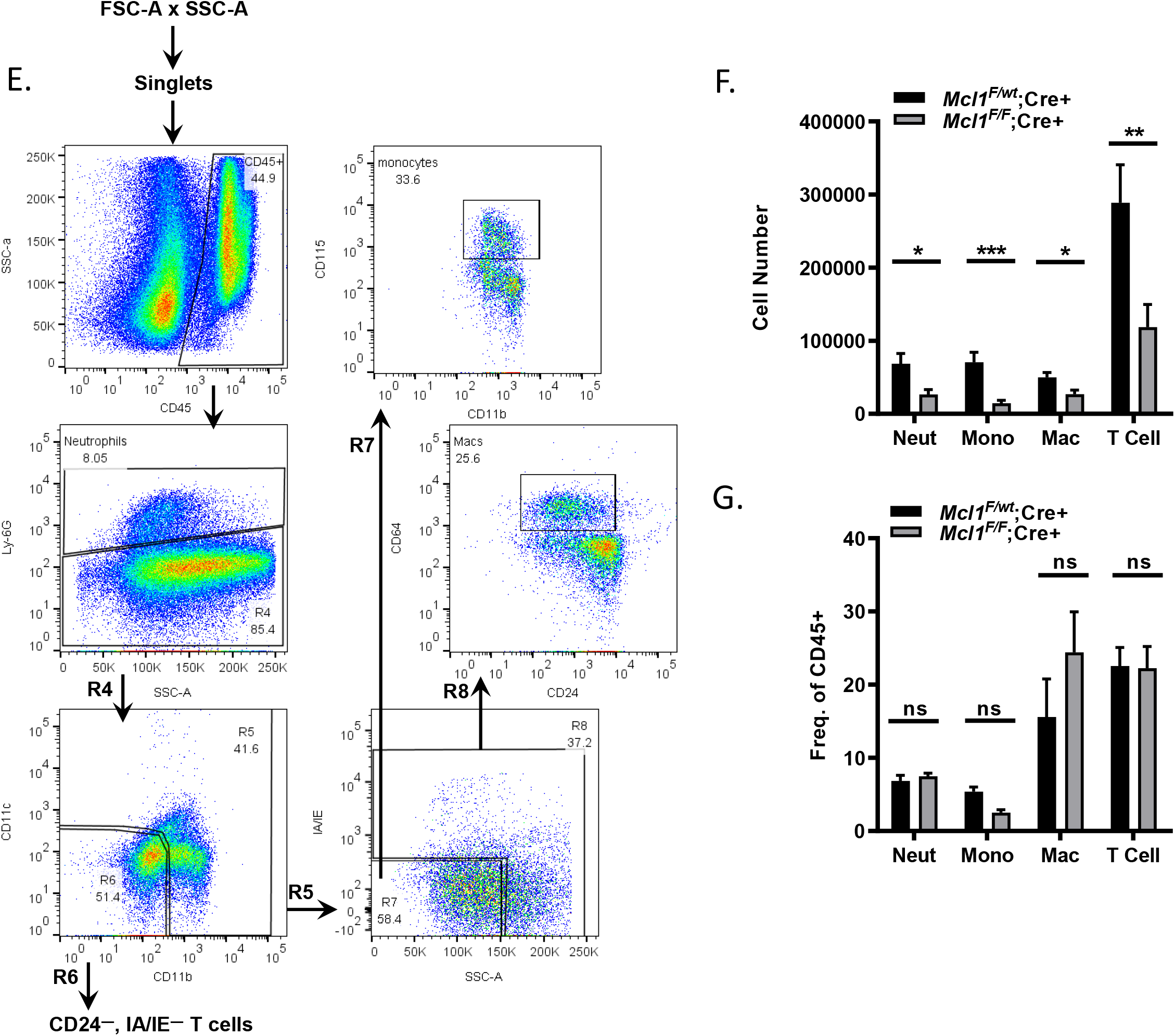
HSC niche is intact in *Mcl1^F/F^* EpoR-Cre+ animals. (A) Gating scheme used to analyze hematopoietic stem cell niche in fetal livers. DAPI staining was used to gate out dead cells. (B) Heat map of fold change in cell number of various progenitor populations. (C) tSNE plots of HSC populations gated on live singlets showing lineage expression (left) or CD45 expression (right) with “WT” representing *Mcl1*^F/wt^; EpoR-Cre+ embryos and “KO” representing *Mcl1^F/F^;* EpoR-Cre+ embryos. (D) Plot of HSC progenitor populations represented as percent lineage negative. (E) Gating scheme used to analyze myeloid compartment in fetal livers. (F) Numbers of cells obtained from fetal livers using gating scheme shown in E. (G) Plot of indicated cell populations, represented as frequency of CD45+. (n=6 litters with at least 3 embryos per genotype for HSC staining; n=5 biological replicates with 3 embryos per group for myeloid staining. Bars represent mean plus or minus SEM, One-way ANOVA with α=0.05. *p<0.05; **p<0.01; ***p<0.001; ns = not significant).

**Supplemental Figure S2.**
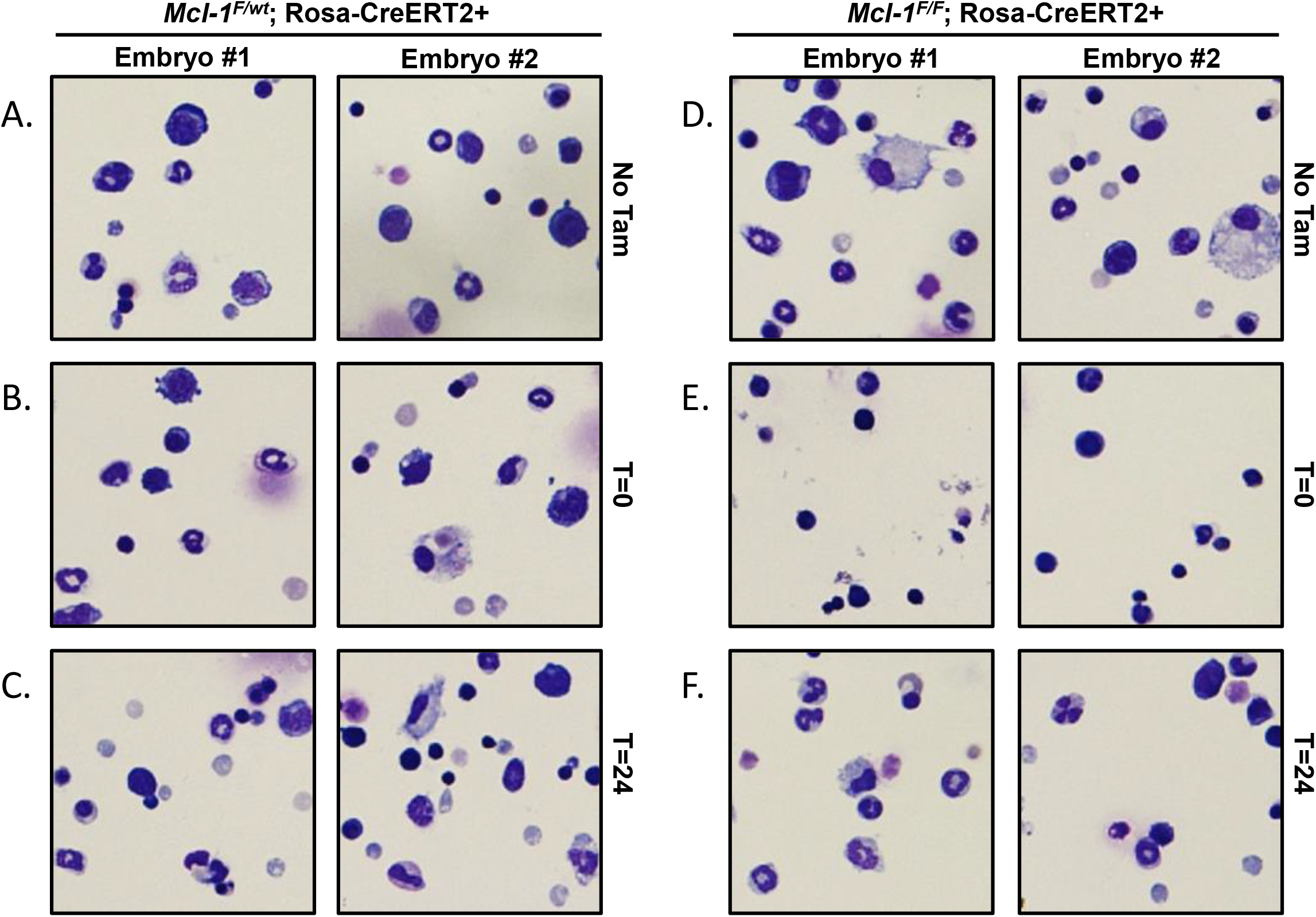
MCL-1 promotes early erythropoiesis but is dispensable during later stages. (A-F) TER119-depleted fetal liver cells from E12.5 *Mcl1^F/F^* Rosa-ERCreT2+ or Mcl1^F/WT^ Rosa-ERCreT2+ littermate controls were cultured for 18 hours in “stem cell media” containing SCF, IL-6, and FLT-3L. (A, D) Media was changed to Epo-containing media without additional stimulus. (B, E) Cells were pulsed with 4-OHT for the first 18 hours of culture (while in stem cell media), then media was changed to Epo-containing media. (C, F) Following 18 hours in stem cell media, cells were placed in mediacontaining 4-OHT, Epo, and other factors (see Methods). Cytospins of 2 different embryos are shown for each culture condition.

**Supplemental Figure S3.**
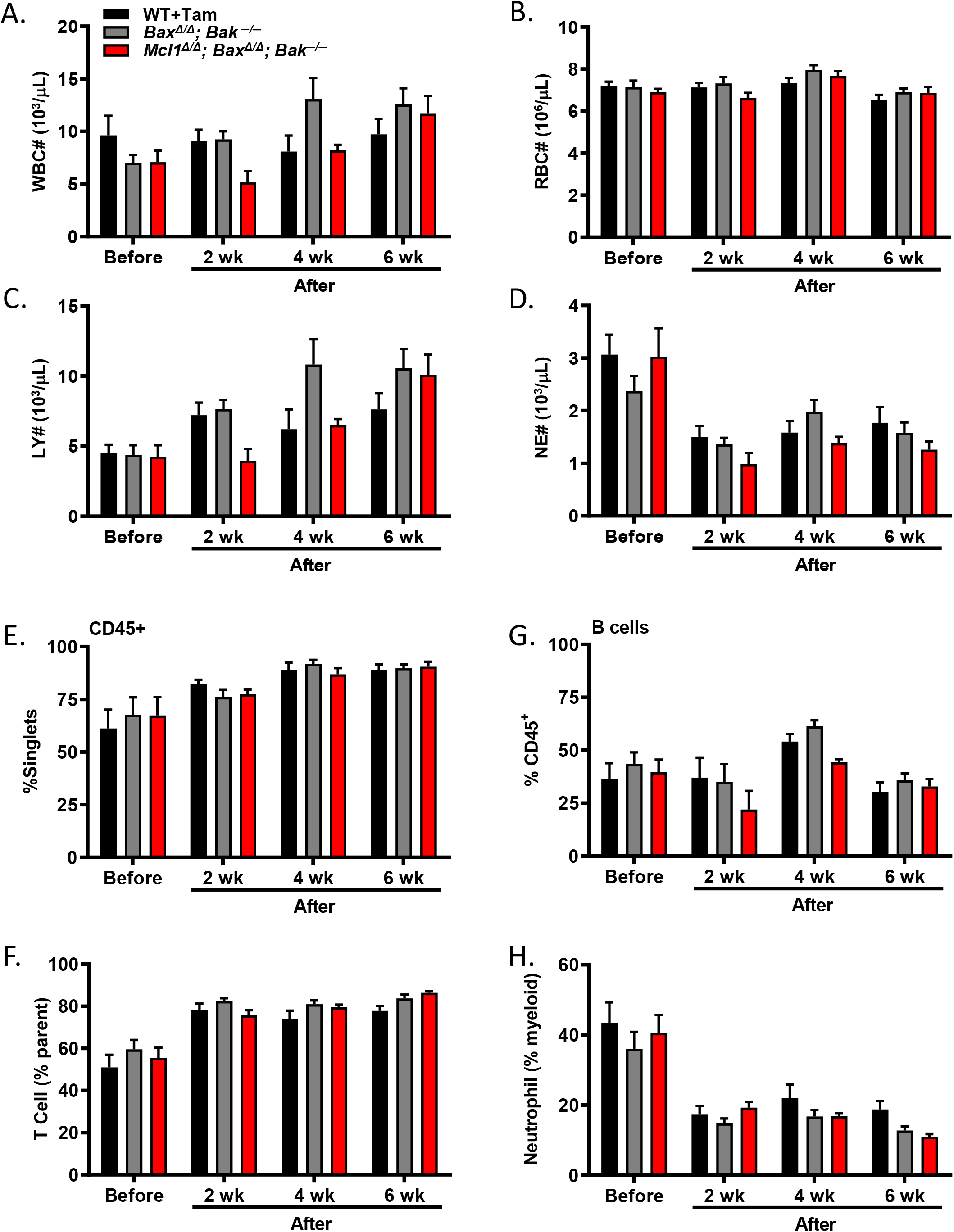
No significant differences in any populations following hematopoietic stem cell transplant and tamoxifen treatment. (A-H) Transplant studies were performed using lethally-irradiated CD45.1+ recipients and three separate groups of CD45.2+ donors: C57BL/6 wildtype, *Bax^F/F^Bak*^−/−^Rosa-ERCreT2, and *Mcl1^F/F^Bax^F/F^BakT^h^* Rosa-ERCreT2 (2 biological replicates with 5-6 recipients per donor group, see Methods). Donor marrow was allowed to engraft for 4 weeks, after which recipient animals received tamoxifen treatment. Blood was taken just prior to tamoxifen treatment (“Before”) then every 2 weeks thereafter. (A-D) Complete blood count (CBC) data indicating white blood cell counts (A), red blood cell counts (B), lymphocyte counts (C), and neutrophil counts (D). (E-H) Flow cytometric analyses of blood indicating CD45+ cells as percent of singlet gate (E); B220+ B cells as percent of CD45+ cells (F); T cells (CD4+ and CD8+) as percent parent population (G); and neutrophils as percent of myeloid (CD11b+) cells (H).

**Supplemental Figure S4.**
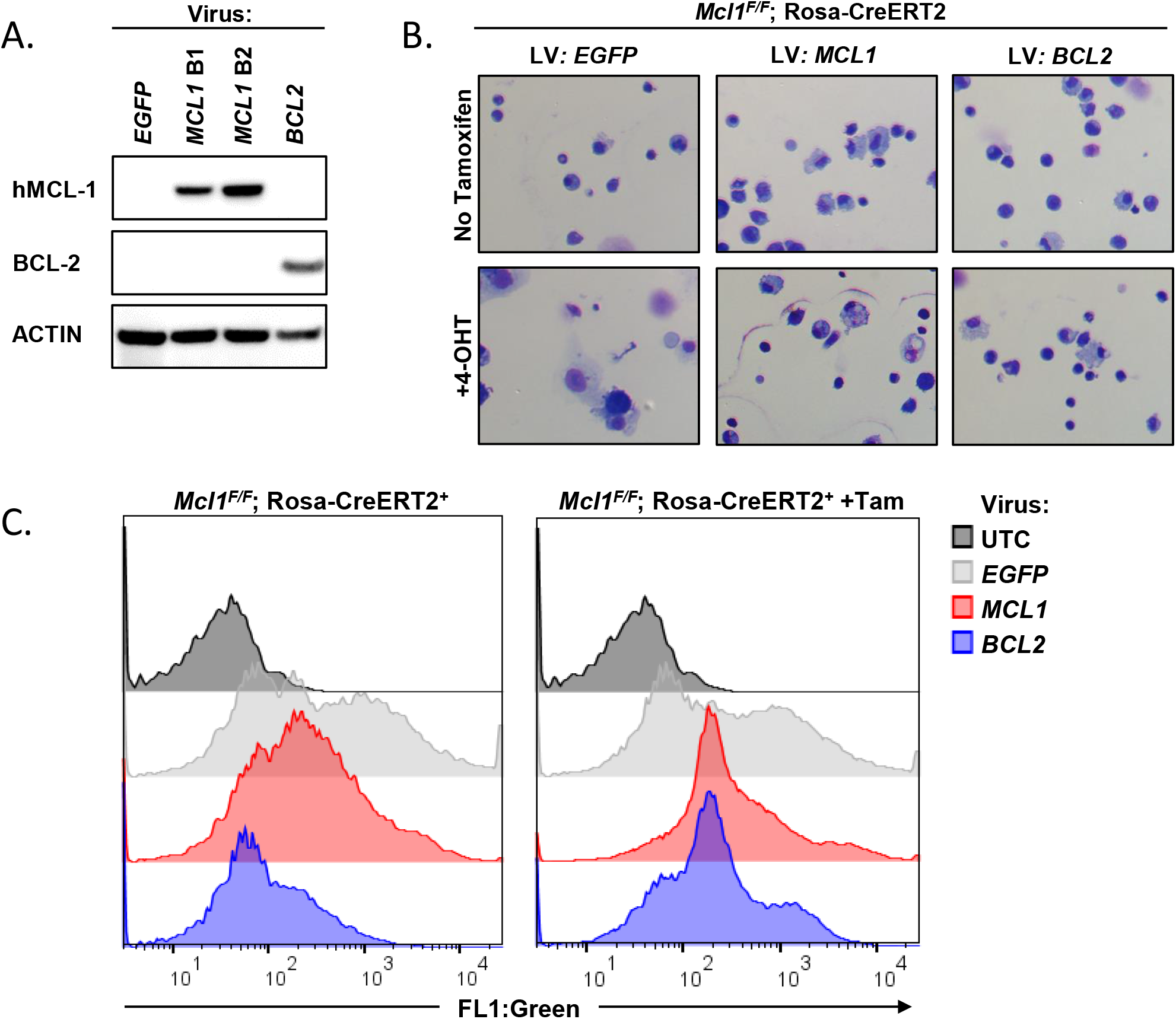
Ectopic expression of human *BCL2* restores erythoid differentiation in the absence of *Mcl1*. (A) Immunoblot from 3T3 cells transduced with the same lentiviruses as used to transduce the fetal liver cells in Figure 4. Lysates were resolved by SDS-PAGE and immunoblotted with the indicated antibodies. Anti-human-specific antibodies were used to detect expression of MCL-1 and BCL-2 and actin serves as loading control. For MCL-1 two independent batches of viral supernatant are presented (B1 and B2). (B) Cytospin preparations from *Mcl1* ‘ Rosa-ERCreT2+ FLC transduced with lentiviral vectors containing *BCL2, MCL1*, of *EGFP* (empty) treated with or without 4-OHT overnight the second day of culture stained with DipQuick staining kit and imaged at 40x. (C) *Mcl1*^F/F^ Rosa-ERCreT2+ fetal liver cells were cultured with the indicated virus for 18 hours, then with or without 4-OHT treatment as in Figure 6. Flow cytometry was performed to assess viral vector expression, indicated by zsGreen fluorescence, before and after 4-OHT treatment. UTC; untransduced control cells.

